# Proteostasis environment shapes higher-order epistasis operating on antibiotic resistance

**DOI:** 10.1101/470971

**Authors:** Rafael F. Guerrero, Samuel V. Scarpino, João V. Rodrigues, Daniel L. Hartl, C. Brandon Ogbunugafor

**Affiliations:** Department of Computer Science, Indiana University; Network Science Institute, Department of Marine and Environmental Sciences, and Department of Physics, Northeastern University; Department of Chemistry and Chemical Biology, Harvard University; Department of Organismic and Evolutionary Biology, Harvard University; Department Ecology and Evolutionary Biology, Brown University; Department of Biology, University of Vermont; Vermont Complex Systems Center, University of Vermont

**Keywords:** epistasis, proteostasis, antibiotic resistance, pleiotropy

## Abstract

Recent studies have shown that higher-order epistasis is ubiquitous and can have large effects on complex traits. Yet, we lack frameworks for understanding how epistatic interactions are influenced by basic aspects of cell physiology. In this study, we assess how protein quality control machinery—a critical component of cell physiology—affects epistasis for different traits related to bacterial resistance to antibiotics. Specifically, we attempt to disentangle the interactions between different protein quality control genetic backgrounds and two sets of mutations: (i) SNPs associated with resistance to antibiotics in an essential bacterial enzyme (dihydrofolate reductase, or DHFR) and (ii) differing DHFR bacterial species-specific amino acid background sequences (*Escherichia coli, Listeria grayi*, and *Chlamydia muridarum*). In doing so, we add nuance to the generic observation that non-linear genetic interactions are widespread and capricious in nature, by proposing a mechanistically-grounded analysis of how proteostasis shapes epistasis. These findings simultaneously fortify and demystify the role of environmental context in modulating higher-order epistasis, with direct implications for evolutionary theory, genetic modification technology, and efforts to manage antimicrobial resistance.

## INTRODUCTION

Vagarious interactions between parcels of genetic information (e.g. mutations, gene variants, gene networks), as captured in phenomena like pleiotropy and epistasis, are widely recognized as a powerful force in crafting the relationship between genotype and phenotype (Cordell, 2002; Remold and Lenski, 2004; Phillips, 2008; Natarajan et al., 2013; Chou et al., 2014; Mackay and Moore, 2014; Sackton and Hartl, 2016; Crona et al., 2017; Otwinowski et al., 2018). Epistasis, informally defined as the “the surprise at the phenotype when mutations are combined, given the constituent mutations’ individual effects” (Weinreich et al., 2013) is now a highly relevant frontier of evolutionary genetics. It casts an uncertainty shadow over many areas of biology aiming to understand or manipulate genetic variation (e.g. GWAS, genetic-modification), as it speaks to unpredictability regarding how phenotypes are related to the genes that are presumed to underlie them.

An especially provocative related phenomenon is “higher-order epistasis.” It offers that parcels of genetic information not only interact in a pairwise fashion (e.g. mutation A interacting non-linearly with mutation B; mutation B interacting non-linearly with mutation C) but potentially in all possible combinations, each with a potentially unique statistical effect (e.g. when the interaction between all 3 mutations—A, B and C—has a quantitative value that cannot be reduced to a combination of independent or pairwise effects) (Weinreich et al., 2013; Poelwijk et al., 2016; Crona et al., 2017; Sailer and Harms, 2017). Statistically, higher-order epistasis is an unwieldy concept because the number of possible interactions can grow exponentially with the number of interacting parcels, which presents both conceptual and computational challenges (as it is a mental challenge to keep track of thousands of potential interactions, and computationally challenging to analyze them using available technology).

Many studies of higher-order epistasis focus on the interactions between suites of SNPs associated with a certain phenotype, engineered in combination or via a library of mutations using high-throughput methods (Ferretti et al., 2016; Poelwijk et al., 2016; Crona et al., 2017; Domingo et al., 2018; Otwinowski et al., 2018; Tamer et al., 2018). Fewer studies specifically dissect the strength and sign of epistatic interactions between SNPs within a gene and particular suites of mutations or gene deletions in other parts of the genome (Williams et al., 2005; Lehner, 2011; Vogwill et al., 2016). Even fewer dissect the impact of physiological contexts on epistasis, a glaring omission when you consider the biochemical and biophysical specifics of the cellular environment in which genes and proteins are made and function.

One particular context that we might predict would shape epistasis within a cell would be that dictated by sets of chaperones and proteases, which have already been demonstrated to have a profound impact on a range of bacterial phenotypes (Gottesman et al., 1997). Prior studies focusing on the chaperonins GroEL/ES and Lon protease have established their centrality in regulating the presence and state of only certain proteins in the cytoplasm (Hartl et al., 2011). And even more recent studies have uncovered how only members of this protein quality control (PQC) system (GroEL/ES and Lon) specifically stabilize different variants of dihydrofolate reductase (Bershtein et al., 2013). The allelic resolution of this proteostasis machinery is a striking finding, and begs the question of how this machinery might frame higher-order epistasis in traits that are controlled by select proteins.

Here, we quantify the magnitude, sign, and order of epistatic effects acting on three mutations within a gene, as influenced by three well-defined proteostasis environments (conferred through the engineering of 3 genotypes of bacteria): wild-type, groEL+, and Δ*lon*. We examine these effects for two related traits that contribute to antibiotic resistance (*IC*_50_ and protein abundance) (Rodrigues et al., 2016), and decompose the impact of proteostasis environment on two classes of potential epistatic interactors: (i) 3 biallelic sites associated with drug resistance in an enzyme target of antibiotics (dihydrofolate reductase or DHFR) and and (ii) 3 different amino acid backgrounds corresponding to species of bacteria (*Escherichia coli, Chlamydia muridarum*, and *Listeria grayi*). We find that the sign and magnitude of interactions among SNPs is highly contingent upon certain genotypic contexts, and observe that epistasis can work differently (qualitatively and quantitatively) across related traits. Importantly, because the biology of the system under study is well understood (e.g. the biophysics of variation in DHFR function, the basis through which PQC machinery regulates proteins), we can surmise on the mechanism underlying certain epistatic interactions both within and between genes. We discuss these findings in light of theory in evolutionary genetics, the study of antibiotic resistance and the challenges facing genetic modification technology.

## METHODS AND MATERIALS

### Strains and phenotypes

Our collection of strains, which are a subset of those originally engineered for the study of the DHFR structure and function by Bershtein et al. (2013), include mutants from three species: *E. coli, L. grayi*, and *C. muridarum*. We measured phenotypic effects of mutations at three sites in the FolA gene enconding DHFR. We encoded the allelic state of a strain using binary notation, 000 corresponding to the ancestor (containing no mutations) and 111 containing all 3 focal mutations, as is common in these types of combinatorial data sets. For simplicity, we refer to individual sites by their position and amino acid change in *E. coli* (even though these can be different in the other two species; see below).

We initially chose *IC*_50_, protein abundance and drugless growth rate as traits of interest. *IC*_50_, a proxy for the ability of an organism to withstand the activity of antibiotics (Trimethoprim in this case), is largely determined by several factors, including abundance and drugless growth rate (Rodrigues et al., 2016).

#### Construction of the PQC mutants

Genes encoding ATP-dependent protease Lon were deleted using homologous recombination enhanced by lambda red, essentially as described (Datsenko and Wanner, 2000). Wild type *E. coli* K12 MG1655 cells were co-transformed with various pFLAG-DHFR mutants and pGro7 plasmid (Takara) expressing groES-groEL under pBAD promoter. Chaperone expression was induced by the addition of 0.2 percent arabinose.

#### Construction of the DHFR mutants

The combinatorially complete set of mutants were constructed for all three species, for the three sites of interest in the FolA gene (*E. coli*, P21L, A26T, and L28R; *C. muridarum*, P23L, E28T, and L30L; *L. grayi*, P21L, A26T, and L28R). These mutations were introduced by Quick-Change Site-Directed Mutagenesis Kit (Stratagene) and cloned into the pFLAG expression vector (Sigma-Aldrich). Each mutagenized plasmid underwent confirmatory sequencing.

#### Measurement of drugless growth rate

Bacterial cultures were grown overnight (37 °C) in M9 minimal medium were normalized to an OD of 0.1 with fresh medium. When appropriate, GroEL overexpression and/or increase in DHFR concentrations were induced by adding arabinose and IPTG immediately after normalization. After additional growth during 5-6 hours a new normalization to an OD = 0.1 was performed before inoculation of 96-well plates (1/5 dilution) containing M9 medium. Growth was quantified by integration of the area under the growth curve.

#### *Measurement of IC*_50_

As with the drugless growth rate, bacteria were grown across a range of concentrations of TMP ranging from (0-2,500 *μ*g/mL), incubated at 37 °C. Absorbance measurements at 600 nm were taken every 30 min for 15 hours. OD readings vs. time were calculated between 0 and 15 hours. *IC*_50_ values were determined from the fit of a logistic equation to plots of growth vs. Trimethoprim concentrations. Reported *IC*_50_ are averaged from at least three replicates.

#### Measurements of intracellular protein abundance

DHFR abundance was measured from the total catalytic activity of the varying alleles in cellular lysate as described in prior work. With regards to protein abundance: how much DHFR is produced by a given cell is the product of many biochemical and biophysical actors. DHFR abundance is an important component of drug resistance, because in order to survive the presence of Trimethoprim (which disrupts the biosynthesis of a folate, a key metabolite; see Supplementary Information), the organism must produce enough DHFR to carry out normal cellular function. Also, because we know that protein quality control machinery, like GroEL and Lon protease, can degrade proteins like DHFR (Bershtein et al., 2013; Rodrigues et al., 2016), then there is a physiological basis for an expectation that these PQC genetic backgrounds would influence protein abundance.

### Statistical analysis

Our approach, a novel application of regularized regression techniques, allows us to measure higher-order epistasis acting across traits and biological scales (both intra- and intergenic). These methods can be used to infer statistical interactions at work in experimental and natural data sets. Notably, this regression approach can be applied to data sets of varying structure, can easily incorporate experimental noise, and can produce results for data sets with missing values (even though the data set in this study is complete). For more discussion on the constraints of other methods commonly used to calculate higher-order epistasis in combinatorial data sets, see the Supplemental Information.

#### Initial exploration

We set out to infer interactions in three bacterial traits: *IC*_50_, DHFR abundance, and bacterial growth rate (total experimental N=232, 360, 252, respectively). For each phenotype, we first fit a general linear model of the form *Y* ~ *S* + *C* + *H*, where *Y* is the phenotype of interest (*IC*_50_, abundance, or growth), *S* is the species fixed factor (with 3 levels), *C* is the PQC context (wild-type, Δ*lon*, and GroEL+) and *H* is a haplotype variable (with 8 levels, coding for the possible combinations of mutations P21L, A26T, and L28R). We tested for the presence of epistasis by fitting alternative models that include the interaction terms *S* × *C, S* × *H, C* × *H* and *S* × *C* × *H*, and choosing the model of best fit based on the Bayesian information criteria (i.e. BIC; a penalty for added regression coefficients proportional to the natural log of the sample size) and a combination of forward and reverse model selection as implemented in the R programming language’s stats package (R Core Team, 2018). After finding significant interaction effects in these initial models for *IC*_50_ and protein abundance, we proceeded to carry out further analyses on these two phenotypes.

#### Regularized regressions (Elastic Net/LASSO)

We tested for epistasis by fitting regularized regressions, which select the set of explanatory variables and estimate their coefficients in a single procedure. Briefly, this is done by including penalties proportional to the value of each coefficient (corresponding to each explanatory variable) in the regression equation. As with other regression procedures (e.g., least-squares), the objective is to minimize this (penalized) equation. In doing so, it finds a balance between small coefficient values and error in the fit of the model. If a variable does not affect the phenotype of interest, its coefficient will be zero. We took non-zero coefficients as evidence that a particular variable, or interaction term, has a significant effect on the phenotype.

We fit these models on standardized phenotypic variables, allowing direct comparisons between the coefficients estimated from different regressions (i.e., units for regression coefficients are standard deviations). Prior to standardization, we log-transformed Abundance and *IC*_50_ values to improve normality. We ran the regression procedures using the glmnet package (Friedman et al., 2010) in R, which carries out an Elastic Net regularization. Specifically, we used the ‘cv.glmnet’ method, which fits models with varying penalty weights (changing the *λ* parameter) and finds the best model by cross-validation (in our case, a leave-one-out approach). To avoid over-fitting, we chose the simplest model that is still within one cross-validated standard error of the best fit model (that is, using *λ_min_* + 1*s.e*.). The Elastic Net method combines linear and quadratic penalties (in *α* and 1 - *α* proportions, respectively) to obtain a sparse set of variables. In the Results section, we present regressions using *α* = 1, which yields fewer non-zero coefficients (and we deem more conservative) and is equivalent to a LASSO approach (Tibshirani, 1996). Regressions using other values of α had little effect on the qualitative patterns (see Data S1 for both data sets: α = 1 and 0.5).

For each phenotype, we fit models at two scales. First, we ran a full model (*Z* ~ *S* × *C* × *P*21*L* × *A*26*T* × *L*28*R*) that included 72 terms: the main effects of five variables (species, PQC context, and the mutations P21L, A26T, and L28R) and all possible interactions. Second, we ran models within PQC-species context (that is, 9 separate models per phenotype), to get a more detailed perspective on how PQC shapes intragenic epistatic interactions. Within each PQC-species context, the model fit was *w* ~ *P*21*L* × *A*26*T* × *L*28*R*, where *w* is the phenotypic value normalized within each group (i.e., using mean and variance of each PQC-species set). The estimated coefficients for these models are summarized in Table S2).

### Script availability

All the scripts for these analyses—written in R (R Core Team, 2018) and using methods in the tidyverse (Wickham, 2017), glmnet (Friedman et al., 2010), and treemapify (Wilkins, 2018) packages—are available in the Supplement.

## RESULTS

We first set out to construct a coarse picture of the experimental data: whole alleles of DHFR, with SNPs in various combination engineered into several background strains, and assayed for three traits relevant to drug resistance. Figure 1 shows how the engineered alleles (the 8 combinatorial mutants) perform with respect to *IC*_50_, protein abundance and drugless growth, across genotypic contexts (species and PQC background). While these first two phenotypes show patterns highly consistent with epistatic interactions at several levels, growth rate shows no significant variance across genotype contexts (confirmed by GLM models and BIC model choice; Figure S1).

**Figure 1.**
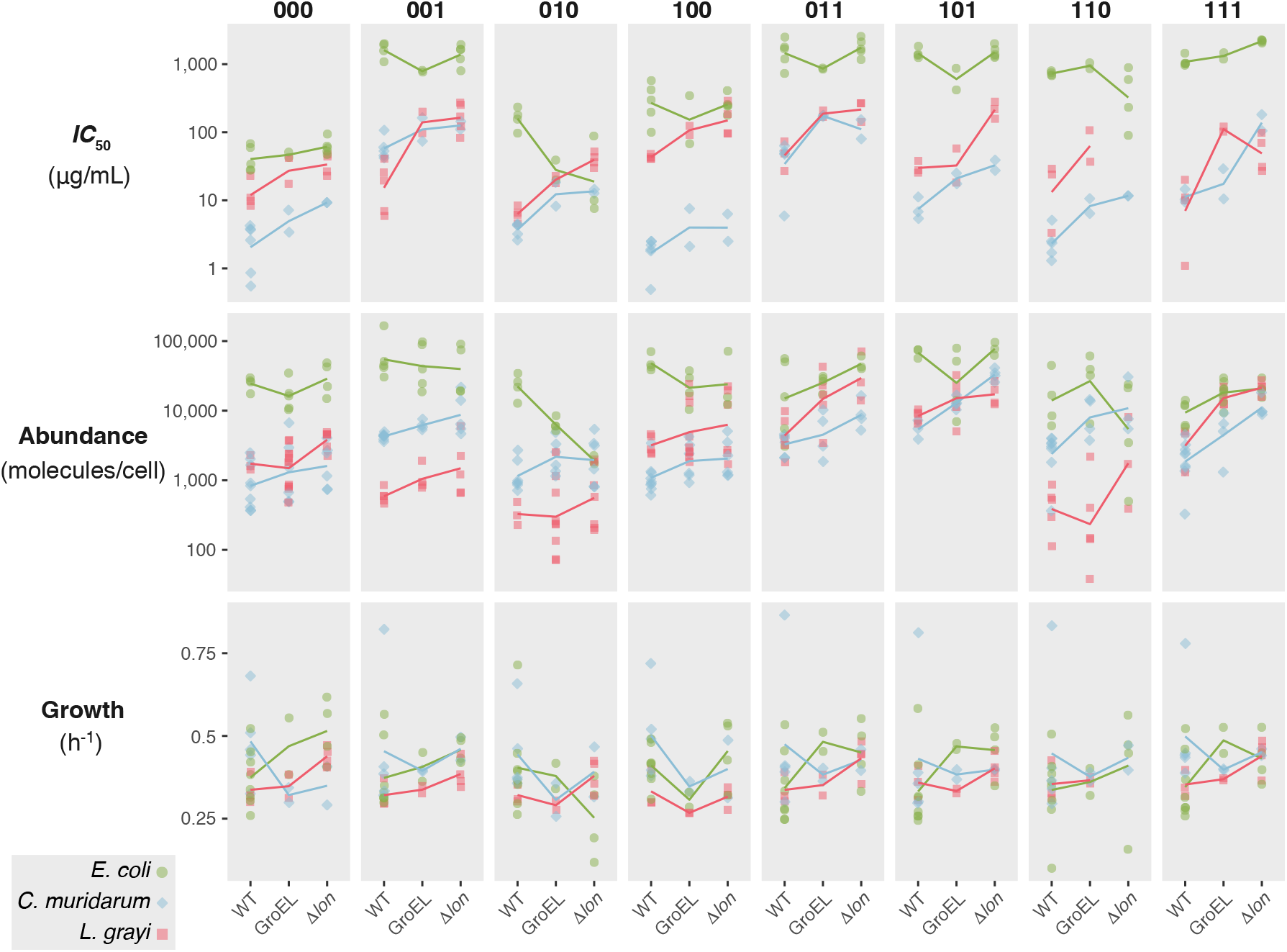
Phenotypic variation of DHFR mutants across proteostasis contexts. Phenotypes depend on protein quality control context (horizontal axis) and species background (*E. coli*, green circles; *C. muridarum*, blue diamonds; *L. grayi*, in red squares). DHFR mutations at three amino acid positions are represented in binary notation (000=wild type, 001= P21L, 010=A26T, 001=L28T, and their respective double and triple mutants). Points are biological replicates; lines are mean values.

Having identified that epistatic interactions are likely to exist in the *IC*_50_ and abundance traits (Figures 1 and S1), we employed a set of regularized regressions to “decompose” the magnitudes, signs and orders of epistatic effects operating at the different scales of genetic parcels represented in this data set (SNPs in DHFR associated with resistance to Trimethoprim, species-specific amino acid background, and PQC mutations). Effect sizes can be found in Tables S1 and S2.

### Decomposition of epistasis for *IC*_50_

The main driver of *IC*_50_ is the species-specific amino acid background (Figure 2A). The *C. muridarum* and *L. grayi* amino acid backgrounds have the largest negative effects in the full LASSO regression for this trait (effect sizes −1.44 and −0.9 respectively). Taken alone, these findings suggest that the species background is an important factor in determining the *IC*_50_ phenotype. Our knowledge of the biology of the system provides us a mechanistically informed interpretation: prior studies demonstrate that the DHFR enzyme in *C. muridarum* is inefficient catalytically, and that *L. grayi* is thermodynamically unstable (Rodrigues et al., 2016). Given that catalysis and thermostability are necessary for an enzyme to carry out its function, that the *C. muridarum* and *L. grayi* amino acid backgrounds have such strong negative effects is unsurprising. We cannot, however, relegate the entirety of main effects to species background: the second-largest effect overall is the presence of the L28R mutation (effect size = 1.22), demonstrating that main effect actors of various kinds can influence the *IC*_50_ phenotype.

**Figure 2.**
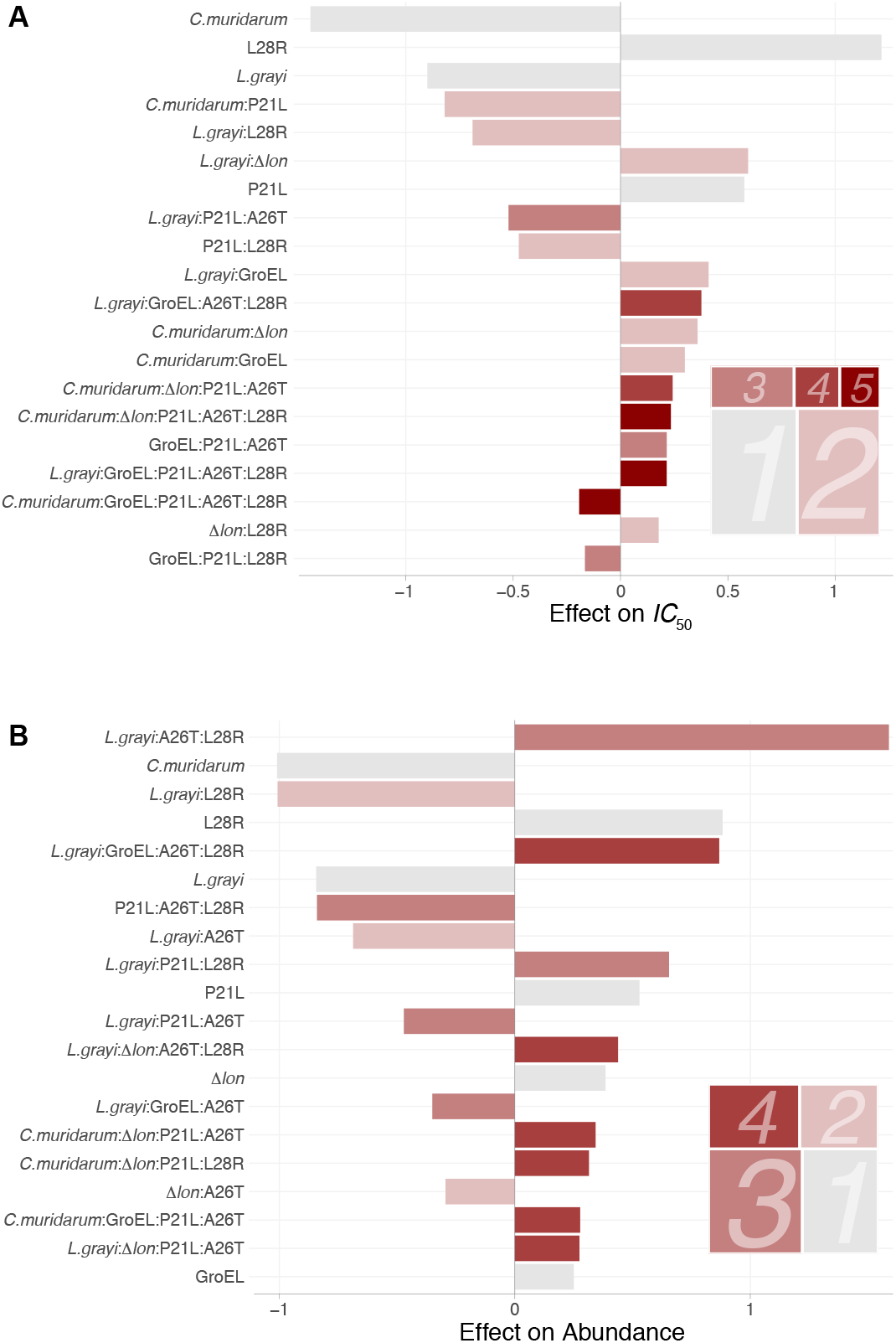
Widespread presence of higher-order interactions for A)*IC*_50_ and B) protein abundance. Distribution of regression coefficients from a LASSO regression allowing interactions across species, protein quality control context, and DHFR mutations for two phenotypes of interest. Bars represent the sign and magnitude of the 20 largest coefficients in the best-fit model. Coefficients are arranged from top to bottom by their magnitude, and their color represents their order (grey for main effects, increasing darkness of red for terms of order 2 through 5). The treemaps in the bottom right corner of each panel represent the sum of all non-zero coefficients by order (the area of each box is the total effect of terms of that order).

Even though main effects define the top three independent drivers of *IC*_50_, higher-order interactions have a larger total effect than main effects on this trait (Figure 2A). Among interactions, the specific patterns are mechanistically diverse: some are between species-specific amino acid background and individual SNPs (e.g, *L. grayi*:L28R, effect size = -0.69), others between species-specific background and PQC environments (*L. grayi*:GroEL+, effect size = 0.41). As with the main effects, several of these findings might be explained by our knowledge of the study system. Though there is a basis for the prediction that the amino acid background of *L. grayi* and the GroEL+ phenotype would interact (the GroEL+ phenotype helps to stabilize the relatively unstable *L. grayi* DHFR enzyme), many of the calculated higher-order interactions cannot be so readily explained, and might serve as the basis of future inquiry. Several plausible mechanistic interpretations are explored in Table 1.

**Table 1.** Possible mechanisms underlying the five largest factors affecting *IC*_50_.

| Effect | Category | Magnitude | Mechanistic interpretation |
| --- | --- | --- | --- |
| *C. muridarum* | Species<br>(main effect) | −1.44 | The *C. muridarum* amino acid background is thermodynamically unstable, more prone to proteolytic degradation and has low catalytic efficiency. Consequently, it has a strong negative effect on the ability to survive in the presence of drug, across all other interacting genetic backgrounds. |
| L28R | SNP<br>(main effect) | 1.22 | The L28R mutation greatly increases both structural stability and the drug inhibition constant ($K_i$), and consequently, helps DHFR perform its enzymatic function in the presence of drug, across genotypic contexts. |
| *L. grayi* | Species<br>(main effect) | −0.90 | The *L. grayi* amino acid background is very thermodynamically unstable and prone to proteolytic degradation. This is partially compensated for by reasonably high catalytic efficiency ($K_{cat}/K_m$), but still has a net negative effect on $IC_{50}$. |
| *C. muridarum*:<br>P21L | Species x SNP<br>(second-order) | −0.82 | The *C. muridarum* amino acid background is inefficient and thermodynamically unstable. The P21L mutation is, however, slightly stabilizing, which diminishes the negative impact of the *C. muridarum* amino acid background. The net effect remains negative however. This result highlights how powerful the *C. muridarum* amino acid background is, in that it can “drag down” the positive effects of certain SNPs. |
| *L. grayi*:L28R | Species x SNP<br>(second-order) | −0.69 | This highlights the non-linear interaction between a powerfully positive SNP (L28R) and the strongly negative main effect *L. grayi* background. That the interaction term is negative highlights that even the stabilizing effects of a positive effect SNP (L28R) cannot compensate for the negative effects of the unstable *L. grayi* amino acid background. |

### Decomposition of epistasis for protein abundance

As with the *IC*_50_, Figure 2B shows that the *C. muridarum* amino acid background has the strongest main effect on protein abundance. This reflects a general pattern of similarity in effects between *IC*_50_ and abundance, which share their top 3 main effect factors: *C. muridarum* (effect size = -1.01), L28R (effect size = 0.88), and *L. grayi* (effect size = -0.84). Interactions, however, appear to play a much larger role in determining protein abundance. We observe several notable patterns, with third-order interactions displaying the largest overall effect, defined by the interaction with the largest single effect (of any) on abundance: *L. grayi*:A26T:L28R (effect size = 1.59). Conspicuously absent from the most important main effects are the PQC backgrounds (GroEL+, Δ*lon*; effect sizes = 0.25 and 0.38, respectively). This suggests that protein quality control machinery is mostly a meaningful actor in determining DHFR abundance in the presence of other genetic parcels. Or rather, only certain SNP and species background combinations seem to be significantly affected by the presence or absence of certain protein quality control variants. Table 2 proposes potential mechanisms that could explain several of these interactions, based on knowledge of the study system. As with *IC*_50_, these proposed mechanisms are speculative, but could be the basis of more detailed inquiry in the future.

**Table 2.**
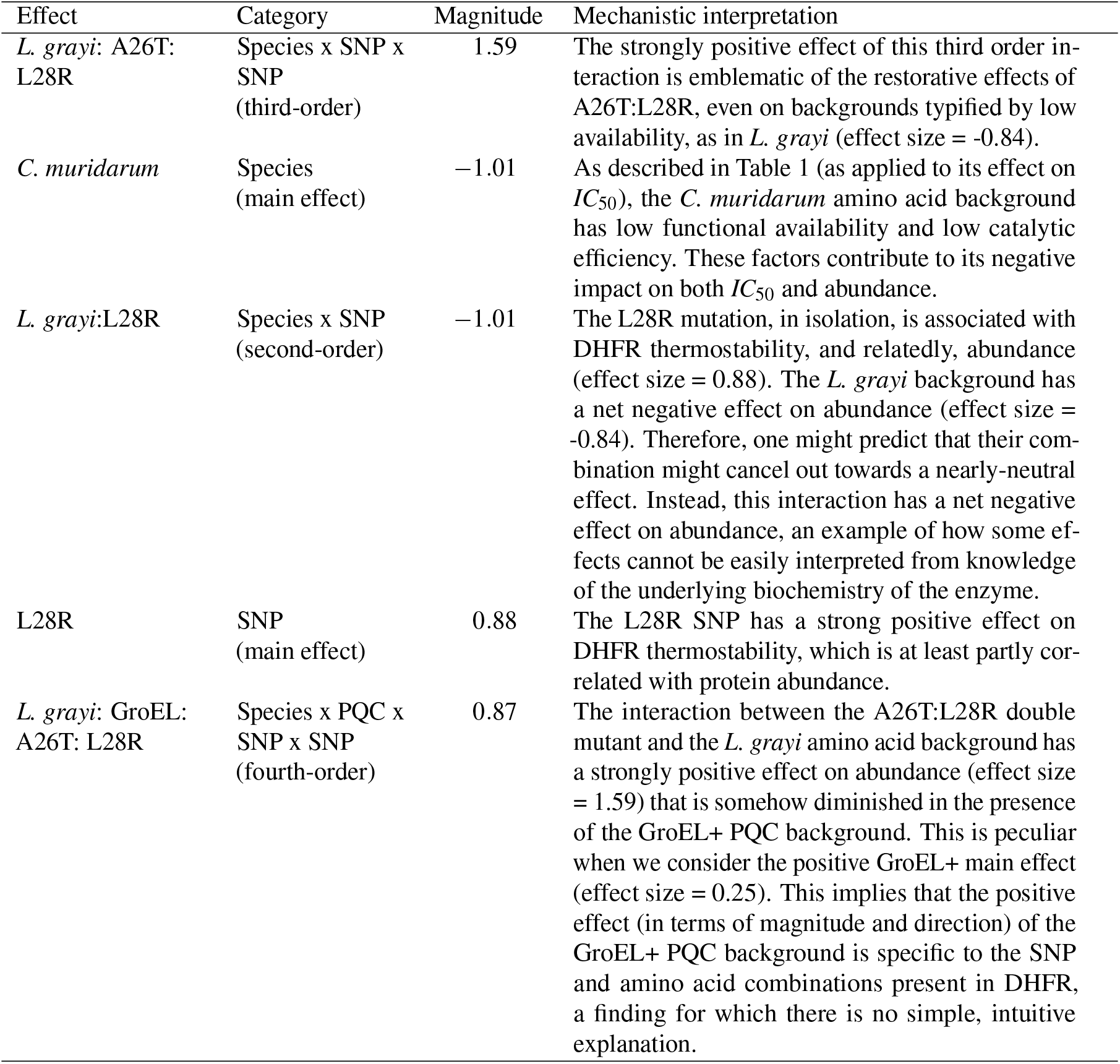
Possible mechanisms underlying the five largest factors affecting DHFR abundance.

| Effect | Category | Magnitude | Mechanistic interpretation |
| --- | --- | --- | --- |
| *L. grayi*: A26T:<br>L28R | Species x SNP x<br>SNP<br>(third-order) | 1.59 | The strongly positive effect of this third order interaction is emblematic of the restorative effects of A26T:L28R, even on backgrounds typified by low availability, as in *L. grayi* (effect size = -0.84). |
| *C. muridarum* | Species<br>(main effect) | -1.01 | As described in Table 1 (as applied to its effect on $IC_{50}$), the *C. muridarum* amino acid background has low functional availability and low catalytic efficiency. These factors contribute to its negative impact on both $IC_{50}$ and abundance. |
| *L. grayi*:L28R | Species x SNP<br>(second-order) | -1.01 | The L28R mutation, in isolation, is associated with DHFR thermostability, and relatedly, abundance (effect size = 0.88). The *L. grayi* background has a net negative effect on abundance (effect size = -0.84). Therefore, one might predict that their combination might cancel out towards a nearly-neutral effect. Instead, this interaction has a net negative effect on abundance, an example of how some effects cannot be easily interpreted from knowledge of the underlying biochemistry of the enzyme. |
| L28R | SNP<br>(main effect) | 0.88 | The L28R SNP has a strong positive effect on DHFR thermostability, which is at least partly correlated with protein abundance. |
| *L. grayi*: GroEL:<br>A26T: L28R | Species x PQC x<br>SNP x SNP<br>(fourth-order) | 0.87 | The interaction between the A26T:L28R double mutant and the *L. grayi* amino acid background has a strongly positive effect on abundance (effect size = 1.59) that is somehow diminished in the presence of the GroEL+ PQC background. This is peculiar when we consider the positive GroEL+ main effect (effect size = 0.25). This implies that the positive effect (in terms of magnitude and direction) of the GroEL+ PQC background is specific to the SNP and amino acid combinations present in DHFR, a finding for which there is no simple, intuitive explanation. |

### *IC*_50_ vs. abundance: correlation and pleiotropy

The determinants of IC50 and protein abundance are similar, but there are meaningful and relevant outliers (Figure 3; *R*^2^ = 0.35, GLM *p* = 10^−7^). The significant relationship between effect sizes estimated from our full models for IC50 and abundance suggests that large-scale patterns of epistasis between these related traits are correlated. This correlation is not surprising: it reflects that these traits are connected at a mechanistic level, since bacteria need to make the enzyme in order to survive the effects of a drug that antagonizes that enzyme. More interesting are, perhaps, the outlier factors: the *L. grayi*:A26T:L28R interaction has a strong effect on abundance (effect size = 1.59) and none on *IC*_50_ (effect size = 0). Similarly, the *L. grayi*:P21L:L28R interaction has a negative effect on *IC*_50_ (effect size = -0.11) and a solidly positive effect on abundance (effect size = 0.66). Thus, at a more detailed level of analysis we observe that individual effects can differ quite substantially, which highlights that certain mutation-interactions can tune related phenotypes in different ways (in both magnitude and sign of effect). The differences in inferred effect sizes suggest that higher-order effects on abundance (P21:A26T:L28R, *L. grayi*:P21L:L28R, *L. grayi*:A26T:L28R) need not translate into downstream effects on *IC*_50_. In other words, we find in these differences some indication of pleiotropy, where mutations (or, in this case, interactions among mutations) display different effects on even functionally-related phenotypes.

**Figure 3.**
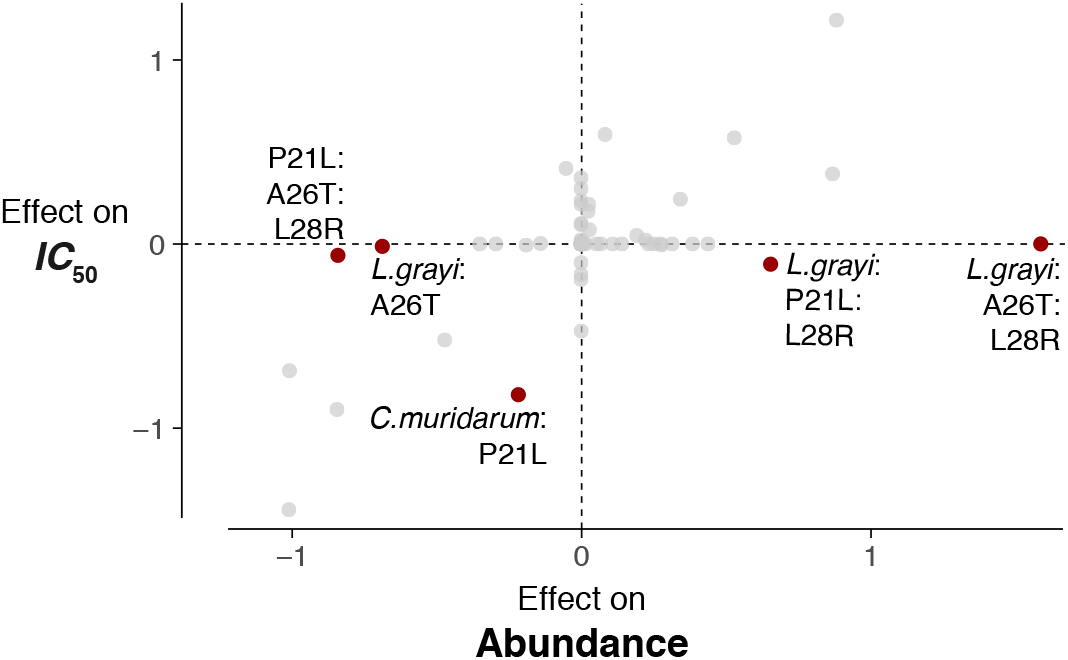
Epistatic effects are correlated between *IC*_50_ and protein abundance traits, with several important higher-order outliers that demonstrate pleiotropic effects. Highlighted are the five terms with largest discrepancies in value between the two phenotypes.

### Epistatic effects of SNPs across PQC contexts

Having conducted analyses aimed at decomposing epistasis across the entire experimental data set (Figures 2 and 3), we employed more granular methods to observe the phenotypic effects of the individual SNPs (P21L, A26T, L28R) in various combination relative to their putative ancestor (genotype 000 in each PQC-species group) as a function of PQC background. The coefficients, inferred by fitting nine separate LASSO models (one per PQC-species background), show considerable variation across PQC backgrounds and are consistent with the notion that PQC background is a direct modulator of epistatic effects. For *IC*_50_, note the especially strong positive effects of the P21L:A26T (E. coli species background; effect size = 1.85) and the A26T:L28R (*L. grayi* species background; effect size = 1.30) pairwise effects in the GroEL+ PQC context. The different PQC backgrounds have markedly different patterns of higher-order epistasis (Figure 4A), with Δ*lon* having notable pairwise interactions across SNP and species amino acid backgrounds.

**Figure 4.**
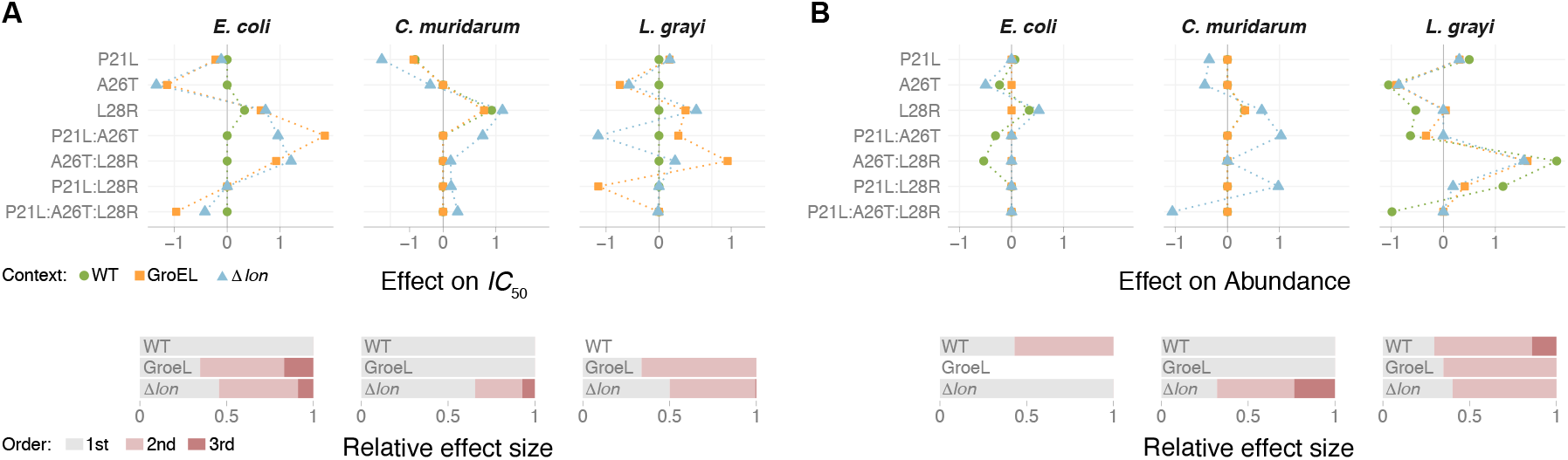
Magnitude, direction and order of epistatic effects between SNPs across PQC-species backgrounds for A) *IC*_50_ and B) DHFR abundance. The estimated effect of single amino acid substitutions (P21L, A26T, and L28R) and their interactions vary across PQC background, indicating higher-order epistasis. Effect sizes were estimated using a LASSO regression within PQC-species background (see Methods and Materials). Dashed lines are drawn for clarity only. The bars at the bottom of each panel summarize the relative contribution of each order (main effects in grey, pair-wise interactions in light red, third-order in dark red) to the total of (absolute) coefficients estimated in each model.

For abundance, PQC background remains a powerful driver of epistatic effects, but in a manner much different than *IC*_50_. In general, epistatic order differed substantially across PQC backgrounds (Figure 4B, bottom panel), with several especially notable effects in Δ*lon*: P21L:A26T and P21:L28R (both in *C. muridarum*; effect sizes = 1.03 and 0.98, respectively), and a third-order interaction P21L:A26T:L28R with a strongly negative effect (also in *C. muridarum*; effect size = -1.06).

## DISCUSSION

In this study, we have attempted to dissect the epistatic interactions (in terms of magnitude, sign, and order) operating across SNPs, species-specific amino acid backgrounds, and protein quality control genetic backgrounds for two phenotypes related to drug resistance in bacteria. Below we discuss the major findings, organized into several subsections. Additional discussion points can be found in the Supplemental Information.

### Higher-order epistasic interactions within and between genes influence two traits related to drug resistance

The results speak to the difficulty in making a priori assumptions about the way that epistasis operates when a system contains potential interactions of different kinds (e.g. intragenic and intergenic). If we assume that physical distance between mutations correlates with the strength of interactions, we might guess that individual SNPs within a gene or local amino acids within the same gene (intragenic epistasis) might interact more readily than parcels between loci (intergenic epistasis). This assumption, however, is not supported by our results: we observed that higher-order interactions involving multiple SNPs and PQC backgrounds (i.e., intergenic interactions) can have important effects on several phenotypes, often as large as intragenic interactions. Discussed in light of modern evolutionary genetics, these results add further color to the debates surrounding the challenges of deconstructing complex phenotypes from effects of individual SNPs, as is often the goal of genome wide association studies. For example, even in circumstances where we are successful in identifying SNPs that are significantly over-represented in a population of individuals with a certain phenotype, interactions between these SNPs and any number of other genetic parcels (perhaps outside of the gene where the candidate SNPs are located) very well may account for most of the variance in the phenotype of interest. That being the case, evolutionary geneticists are justified in being cautious in interpreting the importance of main effect SNPs on complex phenotypes.

### While epistasis patterns are correlated between related traits, several higher-order effects demonstrate pleiotropic behavior

Just as provocative as the observed capriciousness of epistatic interactions is the manner in which these factors influence related traits. Protein abundance affects how a microbe survives the presence of an antibiotic (Trimethoprim in this case) through producing enough DHFR to perform the necessary catalytic functions (which is reflected in the *IC*_50_). Protein abundance has been identified as a component in an quantitative approach used to predict the *IC*_50_ from various biochemical and biophysical parameters (See Supplemental Information). Because of this, we would expect the patterns of epistasis between *IC*_50_ and protein abundance (Figure 4) to be well correlated. At another level of analysis, however, the nature and magnitude of individual effects are different between these traits: several higher-order effects that meaningfully influence protein abundance (both negatively and positively) have almost no effect on *IC*_50_. We might summarize these findings another way: strong overall correlations between epistatic interactions acting on related traits still allows for meaningful differences in the identity and magnitude of individual interactions. When it comes to how certain epistatic interactions manifest, related traits might not be so related at all.

### Patterns of epistasis are broadly affected by PQC environments

We found candidate SNP interactions with large and specific effects on both *IC*_50_ and abundance, but most differed across PQC backgrounds. Though the results in this study have further demonstrated how widespread epistasis can be, we have also identified how there are individual SNPs (or SNP combinations) that influence individual traits while having a minor influence on related ones. And so, despite the prevailing idea that epistasis undermines a simple answer to questions about how complex phenotypes are constructed, our effort to decompose the epistasis in this system has identified SNP/SNP-interactions that could be summarized as being reliable signatures for the phenotypes measured in this study. However, these findings supplement recent studies that emphasize the importance of recipient genome in understanding and predicting the phenotypic effects of transgenic mutations (Vogwill et al., 2016; Wang et al., 2016), as PQC context strongly dictated the consequences of these SNPs.

### Environmental influences on higher-order epistasis: Moving towards mechanistic explanations

An under-nuanced summary of the results of this study might suggest a nihilistic take on modern genetics in light of epistasis, where all roads lead to (hypothetical and sarcastically framed) “epistasis implies that we can never fully decouple the heritable components of a complex trait” or “we can never predict the phenotypic consequences of a given SNP across different genotypic contexts.” These conclusions might be discouraging, especially to those who would prefer that main effects drive the phenotypes of interest (say, in a bio-engineering setting). The data presented here, however, are hardly the only results that would produce such disappointment, as complex traits without higher-order epistasis at work are quickly becoming the exception. That epistasis produces spurious phenotypic effects is an unambiguous theme of the results of this study (reflected most directly in Figures 2 and 4), supporting recent studies that affirm the presence of higher-order epistasis across a wide breadth of phenotypes, in many organisms.

Moreover, we argue that such broad summaries of epistasis patterns are unnecessary, as our analysis allows us to discuss epistasis at a greater (and more useful) level of detail. We specifically demonstrate how individual components of a critical physiological determinant (PQC) environment shape how epistasis manifests in a single protein, across phenotypes. Note that, in prior studies, GroEL+ and Δ*lon* were demonstrated to have similar effects on DHFR mutations. In this study, their respective cytoplasmic environments shaped higher-order interactions differently (however the magntidude) across different organismal traits. For example, the results suggest where to start if we ever wanted to tune the phenotypes in this study in a certain direction. We found that the L28R main effect has a positive influence on *IC*_50_ in many contexts, and that the A26T:L28R combination powerfully influences DHFR abundance in the *L. grayi* background.

Lastly, our approach does more than simply resolve how epistatic interactions drive a set of phenotypes. These results also offer a small step towards what might be the future of the study of epistasis, where statistical methods reveal potential mechanisms or generate testable hypotheses for how parcels of genetic information interact in constructing complex phenotypes. This perspective will be necessary if true genetic-modification (as driven by CRISPR or other methods) will ever become commonplace. At some point we will need to know what to expect when we engineer a given mutation into a given background: how that mutation interacts with other parcels (across the genome), how we might finely tune such interactions, or if we should bother trying at all.

## ACKNOWLEDGMENTS

The authors would like to thank S. Almagro-Moreno, M. Eppstein, C. Marx, and D. Weinreich for helpful discussions. CBO acknowledges funding support from NSF RII Track-2 FEC (Award Number: 1736253), “Using Biophysical Protein Models to Map Genetic Variation to Phenotypes.” RFG is supported by the Precision Health Initiative at Indiana University.

## SUPPLEMENTAL INFORMATION

### Summary

Below find additional treatments of sub-topics which may be relevant to material offered in main text. These include a supplemental discussion, methods and results. They also include supplemental figures and tables.

### Study limitations

As with any study making a general claim about an important problem (epistasis in this case), there is the ubiquitous potential critique about the possibility that these results “do not generalize.” We remind the authors of such criticism that the study system focused on traits related to antibiotic resistance, a phenotype with biomedical implications. That being the case, even if the methods and results were only relevant to the problem of antibacterial resistance to antifolate compounds (and did not generalize further), we would consider the findings to be relevant for several scientific and biomedical communities. We are, however, confident that the methods and results are reflective of phenomena present in complex traits across the biosphere.

### Further notes on motivations for epistatic decomposition

Several studies that measure epistasis utilize data sets where multiple mutations are constructed in all possible combinations, often in the guise of a graph called a fitness (or adaptive) landscape (Greene and Crona, 2014; Ferretti et al., 2016; Ogbunugafor et al., 2016; Sailer and Harms, 2017; Weinreich et al., 2018). For data sets where variation at sites of interest is biallelic, these combinatorial sets are composed of 2^*L*^ mutations, where *L* is the number of different loci being examined. The mutations that compose the combinatorial set might have originated from experimental evolution (Chou et al., 2014; Toprak et al., 2012) or from field surveys (Projecto-Garcia et al., 2013; Domyan et al., 2014; Natarajan et al., 2018). Regardless of their source, several methods have been introduced to detect the presence of higher-order interactions between mutations in combinatorial data sets. One notable method involves the Fourier-Walsh transformation to generate terms corresponding to epistatic interactions between biallelic sites in a fitness graph (Weinreich et al., 2013; Poelwijk et al., 2016; Weinreich et al., 2018). The benefits of this approach include the transparency and relative ease of the calculation: given a complete fitness graph, one can get a good estimation of how much higher-order epistasis is operating. These methods, while effective, are defined by several constraints: the phenotype values in the graph used to calculate these epistatic terms do not effectively incorporate experimental noise into their calculations. In addition, these methods are mathematically confined to combinatorial sets of SNPs that are biallelic. Lastly, these methods do not apply to data sets where there are missing values, a characteristic of many experimental and natural data sets. Even more modern methods of measuring genetic interactions based on the fitness rank-order of alleles can estimate epistasis in partial data sets (Crona et al., 2017). These methods have, to the best of our knowledge, so far been applied to fitness graphs composed of biallelic loci.

As discussed in the Methods and Materials section, the epistatic decomposition methods utilized in this study have no such constraints, as they incorporate experimental noise, do not require biallelic loci, and can accommodate missing data. The relaxing of the biallelic loci constraint is especially important for this study: while the individual SNP loci in the data set can be characterized as biallelic (P21L, A26T, L28R), the species context (Escherichia coli, Chlamydia muridarum, and *Listeria grayi*) and protein quality control genetic background (wild-type, groEL+, and Δ*lon*) are each composed of three variants per locus.

### 0.1 Notes on the choice of traits for study: drugless growth, *IC*_50_, and intracellular abundance

The biology underlying how the measured traits (drugless growth, *IC*_50_, intracellular abundance) relate to drug resistance is well-studied and reasonably intuitive. Drugless growth rate is synonymous to fitness of an organism. In order to be resistant to Trimethoprim, a given microbe must demonstrate some baseline ability to grow. In this system, we expect drugless growth to be lower and less variant across genotypic contexts, indicative of a trait with relatively little higher-order epistasis. This expectation comes from our knowledge of the biology of the system: plasmids were used to express the DHFR mutants in the background bacterial strain in order to measure abundance, *IC*_50_ and drugless growth. In almost all strains, the simple presence of the plasmid was burdensome to the background strain, almost independent of which species of DHFR was being expressed, or what the PQC genetic background was. Consequently, the drugless growth trait provides something analogous to a negative control, a trait that should be relatively bereft of higher-order epistasis, as all bacterial strains carrying plasmid had a similar low growth rate.

### Notes on the biochemistry and biophysics of the study system

Prior studies have established that the deleterious effect of destabilizing DHFR mutations can be alleviated by the action of the protein quality control(PQC) machinery (Bershtein et al., 2013). Specifically, GroEL/ES chaperonins and Lon proteases were shown to be major modulators of the total intracellular DHFR abundance, acting upon partially unfolded protein intermediates to either promote folding or proteolytic degradation, respectively. The impact of PQC background on fitness is particularly relevant in cases where drug-resistance DHFR mutations are associated with stability trade-offs (Rodrigues et al., 2016) and in scenarios of horizontal gene transfer (Bershtein et al., 2015).

Though the *IC*_50_ values utilized in this study are laboratory derived, prior studies have identified relationships between *IC*_50_ and biochemical and biophysical parameters. Rodrigues et al. Rodrigues et al. (2016) described such an analytical expression:

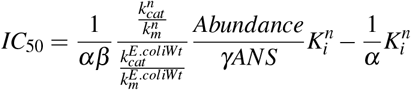

We do not use the above equation in this study, and consequently, are providing it in the supplemental information only to highlight that a mathematical relationship has been proposed that links these traits analytically. Mechanistically, we can most simplistically summarize their relationship this way: in order for a population of bacteria to grow in the presence of trimethoprim (which *IC*_50_ is a presumptive measure of), they must be functional cells that can growth without drug and must produce enough DHFR (the target of Trimethoprim) such that the normal metabolic functions of DHFR are performed. If only small amounts of DHFR are produced, then we can expect low amounts of drug to sufficiently limit growth (low *IC*_50_).

### Regarding antifolates and the evolution of resistance

The study focused on dihydrofolate reductase, an essential enzyme and target of antibiotics. Though the focus of the study was more general (about resolving epistastic effects across genotypic contexts), the specific biological problem of antifolate resistance did warrant a more detailed examination, which we provide here. Antifolates are used clinically as treatments for a wide range of diseases, ranging from bacteria, to protozoal diseases and as anticancer agents (Bershtein et al., 2015; Schnell et al., 2004; Kompis et al., 2005; Liu et al., 2013). These compounds interfere with one of two steps in the *de novo* biosynthetic pathway of tetrahydrofolate (THF), essential for the production of purines and of several amino acids. The genetic basis for antifolate resistance evolution in bacteria lies in a small number of missense mutations in several genes, one of which is dihydrofolate reductase (DHFR). Previous studies had identified that three mutations (A26T, P21L, L28R) that are often found in present various combinations and have an effect on trimethoprim resistance (an antifolate) in *E. coli* (Toprak et al., 2012).

### Regarding the implications of the results for the study of antibiotic resistance

We should very briefly highlight the results in light of their implications for the study of drug resistance. As previously described, the study system was bacterial DHFRs, the protein target of antifolate drugs. While we might call these drugs “antifolates,” we should be very clear about how genotypic contexts influence the evolution of drug resistance. Consequently, for future efforts at “resistance management,” we should be clear about what contextual details influence the phenotypic consequences of resistance-associated SNPs before we fully conclude how a given set of SNPs drives resistance evolution in nature. Direct questions about what these data say about the evolution of drug resistance are the object of current inquiry from individuals involved in this study.

**Supplemental Table 1.** Epistastic decomposition: Regression effect sizes by order for *IC*_50_, protein abundance, and drugless growth, for *α* = 0.5 and 1.0.

**Supplemental Table 2.** Transgenic SNP analyses. These are the data displayed in Figure S2, that demonstrate the phenotypic effects of individual SNP and SNP-combinations.

**Supplemental Table 3.** Biophysical properties of the mutants as measured in prior studies (Rodrigues et al., 2016). We supply them here because they are the basis for speculations on the mechanisms underlying some of the epistatic interactions measures in this study (as discussed in Tables 1 and 2).

**Supplemental data and code.** Data and scripts used in this study can be found at the following location: https://github.com/guerreror/dhfr

**Figure S1.**
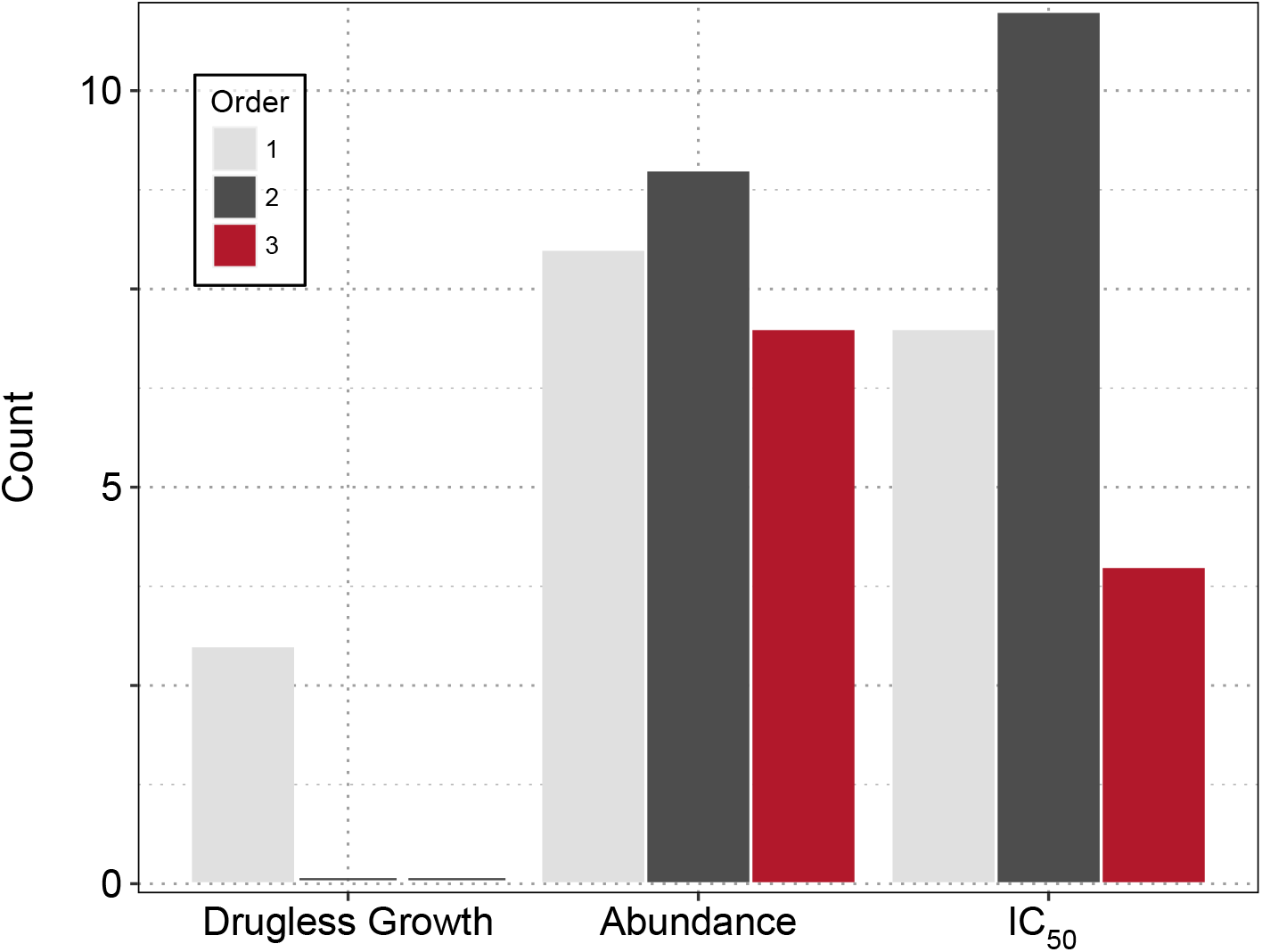
Based on the Bayesian inference criteria (BIC), drugless growth rate has no higher-order interactions, and very few significant main effect drivers. This is in stark contrast to the *IC*_50_ and abundance phenotypes, both which of contain several higher-order interactions.

